# Symmetrically substituted dichlorophenes inhibit *N*-acyl-phosphatidylethanolamine phospholipase D

**DOI:** 10.1101/2020.03.05.979567

**Authors:** Geetika Aggarwal, Jonah E. Zarrow, Zahra Mashhadi, C. Robb Flynn, Paige Vinson, C. David Weaver, Sean S. Davies

## Abstract

*N*-acyl-phosphatidylethanolamine phospholipase D (NAPE-PLD) (EC 3.1.4.4) catalyzes the final step in the biosynthesis of *N*-acyl-ethanolamides (NAEs). Reduced NAPE-PLD expression and activity may contribute to obesity and inflammation, but a major obstacle to elucidating the role of NAPE-PLD and NAE biosynthesis in various physiological processes has been the lack of effective NAPE-PLD inhibitors. The endogenous bile acid lithocholic acid (LCA) inhibits NAPE-PLD activity (IC_50_ 68 μM) but LCA is also a highly potent ligand for TGR5 (EC_50_ 0.52 μM). Recently, the first selective small molecule inhibitor of NAPE-PLD, ARN19874, was reported (IC_50_ 34 μM). To identify more potent inhibitors of NAPE-PLD, we screened compounds using a quenched fluorescent NAPE analog, PED-A1, as a substrate for recombinant mouse NAPE-PLD. Screened compounds included a panel of bile acids as well as a library of experimental compounds (the Spectrum Collection). Muricholic acids and several other bile acids inhibited NAPE-PLD with potency similar to LCA. Fourteen potent NAPE-PLD inhibitors were identified in the Spectrum Collection, with the two most potent (IC_50_ ~2 μM) being symmetrically substituted dichlorophenes: hexachlorophene and bithionol. Structure activity relationship assays using additional substituted dichlorophenes identified key moieties needed for NAPE-PLD inhibition. Both hexachlorophene and bithionol showed significant selectivity for NAPE-PLD compared to non-target lipase activities such as *S. chromofuscus* PLD activity or serum lipase activity. Both also effectively inhibited NAPE-PLD activity in cultured HEK293 cells.

## Introduction

*N*-acyl-ethanolamide (NAE) biosynthesis and signaling appear to play important roles in regulating energy balance, inflammation, stress responses, and addiction (1–6). Saturated and monounsaturated NAEs like *N*-palmitoyl-ethanolamide (C16:0NAE) and *N*-oleoyl-ethanolamide (C18:1NAE) act on GPR55 and GPR119, respectively, as well as on PPARα (7–13). Pharmacological administration of these compounds enhances the resolution of inflammation, induces satiety, and protects against the development of obesity on high fat diet (14–17). In contrast, *N*-arachidonoyl-ethanolamide (C20:4NAE, anandamide) is a polyunsaturated NAE that acts at endocannabinoid receptors (CB1 and CB2) to exert pleotropic effects on food intake, anxiety, nociception, inflammation, locomotion, and memory (7,18).

Even with recent advances, much about the regulation of NAE levels and their contributions to biological processes remains poorly understood. The first step of NAE biosynthesis requires the transfer of the appropriate *O*-acyl chain from phosphatidylcholine (PC) to the ethanolamine headgroup of a phosphatidylethanolamine (PE) to form *N*-acyl-phosphatidylethanolamines (NAPEs) (19). Five calcium-independent PE *N*-acyltransferases (PLAAT1–5) and one calcium dependent PE *N*-acyltransferase (PLA2G4e) that catalyze this transfer have been identified in humans (19–20). The second step of NAE biosynthesis, conversion of NAPE to NAEs, can be directly catalyzed by NAPE phospholipase D, a member of the zinc metallo-β-lactamase family (Figure 1) (21–23). NAPE-PLD homologs are found in mammals, reptiles, worms, and yeast, suggesting that it has highly conserved functions in normal physiology (12,21,24–25). Surprisingly, global deletion of the *Nape-pld* in mice from birth causes marked changes in multiple lipid pathways yet only partially reduces NAE levels in most tissues (26,27–29). Several alternative pathways for the conversion of NAPE to NAE have now been identified (Figure 1) (24,30–31). Despite the identification of these enzymes, how biosynthesis of individual NAEs is controlled remains unclear. For instance, re-feeding of rodents leads to an increase in jejunal OEA levels, but also leads to a decrease in jejunal AEA levels (32), despite the notion that both NAEs theoretically share the same biosynthetic machinery. Modulating the NAE-biosynthetic enzymes with appropriate tool compounds should help elucidate the contribution of individual enzymes to the biosynthesis of individual NAEs under various physiological and pathophysiological conditions and could lead to the development of effective therapies for a range of clinical conditions including obesity, inflammation, chronic pain, and addiction (33–34).

**Figure 1.**
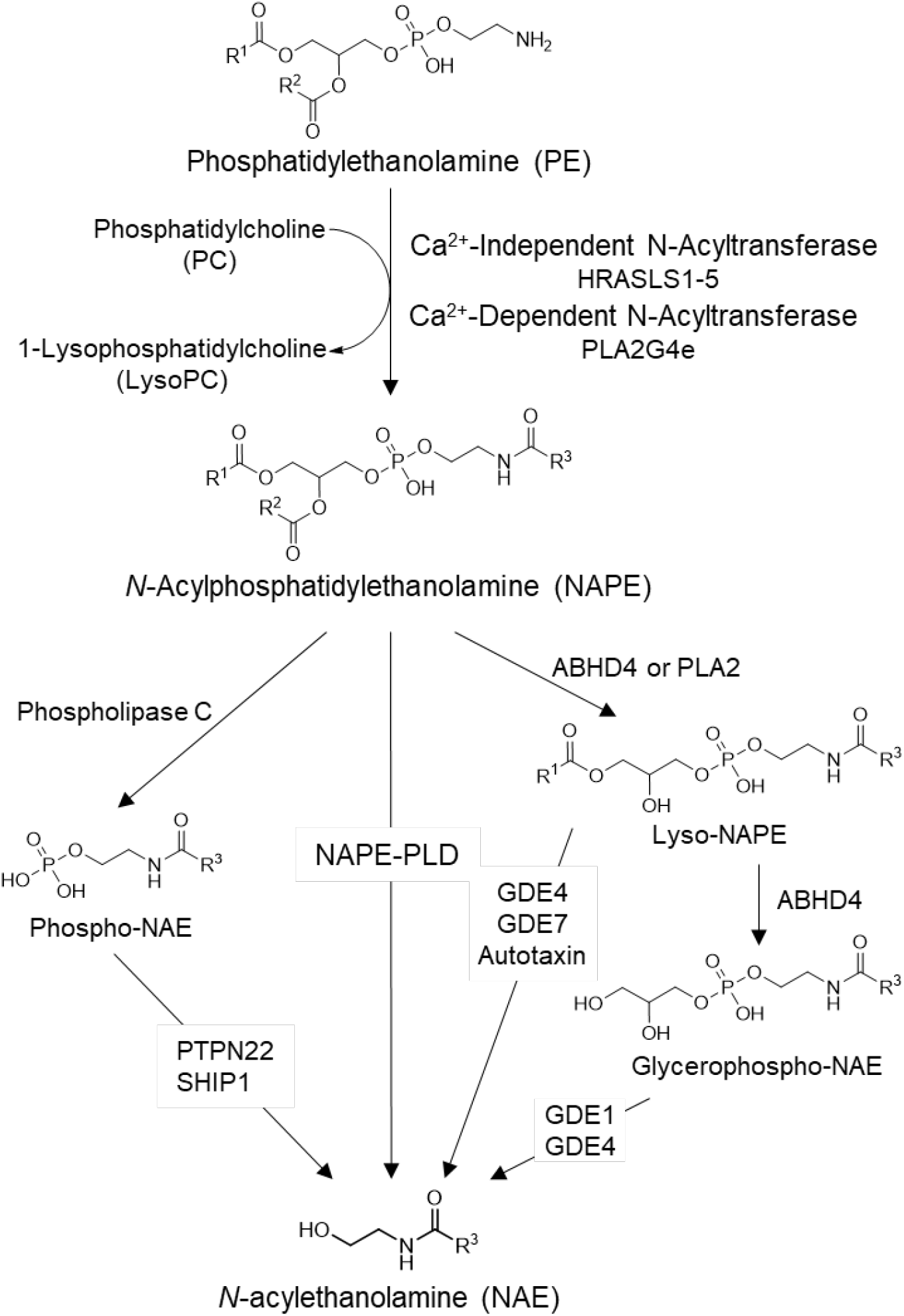
Pathways of *N*-acylethanolamide (NAE) biosynthesis from NAPE. a/b-hydrolase 4 (ABH4), Glycerophosphodiester phosphodiesterase (GDE) −1, −4, −7, *H*-ras like suppressor protein (HRASLS) 1-5, Phospholipase A2 (PLA2), PLA2 group 1Ve (PLA2G4e), Protein tyrosine phosphatase non-receptor type 22 (PTPN22), R1-3, acyl chain substituents, SH-2 containing inositol 5’ polyphosphatase 1 (SHIP-1).

There is currently a lack of specific inhibitors or activators to directly assess the contribution of NAPE-PLD to NAE biosynthesis under various conditions. One endogenous bile acid, lithocholic acid (LCA), has been reported to inhibit NAPE-PLD (IC_50_ 68 μM) (35). Although its bioavailability and relatively low toxicity allow LCA to be administered in vivo, LCA is also a potent ligand for the bile acid receptor, Tgr5, with an EC_50_ 0.53 μM (36) so it cannot be used as a selective inhibitor of NAPE-PLD. More recently, ARN19874 was reported to be a selective NAPE-PLD inhibitor, with an IC_50_ of 34 μM (37). The absorbance, distribution, metabolism, and excretion (ADME) properties of this compound and whether it has efficacy in vivo have not been reported. We therefore performed high-throughput screening (HTS) for modulators of recombinant mouse Nape-pld using a small chemical library consisting of a collection of U.S. and European drugs with known ADME characteristics (the Spectrum Collection), and identified 14 Nape-pld inhibitors with IC_50_ <20 μM. The two most potent compounds were symmetrically substituted dichlorophenes which showed at least 75-fold specificity towards Nape-pld over other lipases tested, and effectively inhibited NAPE-PLD in HEK293 cells.

## Results and discussion

An optimal HTS assay for changes in NAPE-PLD activity requires high reproducibility for replicate samples and sufficient dynamic range to reliably detect activators and inhibitors. Fluorogenic lipid substrates have been previously used successfully in HTS assays for modulators of lipase activity (38–39). PED-A1 is a quenched fluorogenic NAPE analog that has previously been used to measure phospholipase A1 (PLA1) activity in vitro and in tissue samples (40). To test if PED-A1 could be used as a substrate for NAPE-PLD, we incubated PED-A1 with hexa-histidine tagged recombinant mouse Nape-pld and measured the development of fluorescence. Incubation of PED-A1 with active Nape-pld resulted in a rapid rise in fluorescence while incubation with Nape-pld that had been inactivated by boiling did not (See Figure S1A in Supporting Information). The KM of recombinant Nape-pld for PED-A1 (3.0±0.5 μM) was determined by varying the PED-A1 concentration incubated with 0.1 μM Nape-pld (See Figure S1B in Supporting Information). To ensure that the assay would tolerate small variations in buffer composition, we determined the concentration range of N-octyl-β-D-glucoside (NOG, used to stabilize Nape-pld) and DMSO (used as vehicle for PED-A1 and for screening compounds) where Nape-pld activity did not differ by more than 10% from maximal activity. For NOG, this was 0.2–0.6% (w/v) and for DMSO this was 1.0–3.7% (v/v) (data not shown). Based on these results, a final concentration of 0.4% NOG (w/v) and 1.6% DMSO (v/v) was used in all subsequent assays. To assess the assay’s ability to reproducibly distinguish between normal and inhibited signals across replicated wells in 384-well plate, 100 μM LCA or vehicle were added in checkerboard fashion to a 384-well plate and initial rate of fluorescence change (t = 30–100s) and steady state fluorescence (t = 419–420s) determined (See Figure S2 in Supporting Information). The Z’ factor calculated for LCA (0.45–0.47) demonstrated that the signal windows were sufficiently separated to observe hits in the screen, regardless of plate position.

LCA is currently the only known endogenous inhibitor of NAPE-PLD (35). There are a variety of primary and secondary bile acids besides LCA and these differ from each other in respect to the number, position, and stereochemistry of their hydroxyl groups as well as their conjugated moieties. To determine if other bile acids serve as endogenous inhibitors of Nape-pld, we screened a panel of 19 primary or secondary bile acids (50 μM each) (Figure 2). The LCA conjugates glycolithocholic and taurolithocholic acid were slightly less potent Nape-pld inhibitors than LCA, which suggests that there is some flexibility in the length of the negatively charged moiety needed for inhibitory interactions with Nape-pld. Taurine conjugates of deoxycholic acid (tauroursodeoxycholic acid and taurohyodeoxycholic acid) were nearly as potent as LCA, while their non-conjugated forms (deoxycholic acid and ursocholic acid) had no inhibitory effects. Finally, several muricholic acids (α-, β-, and tauro-α-muricholic acids) were slightly less potent than LCA. These results identify a number of new endogenous inhibitors of Nape-pld that may be relevant to Nape-pld activity in the intestinal tract where bile acid concentrations are high.

**Figure 2.**
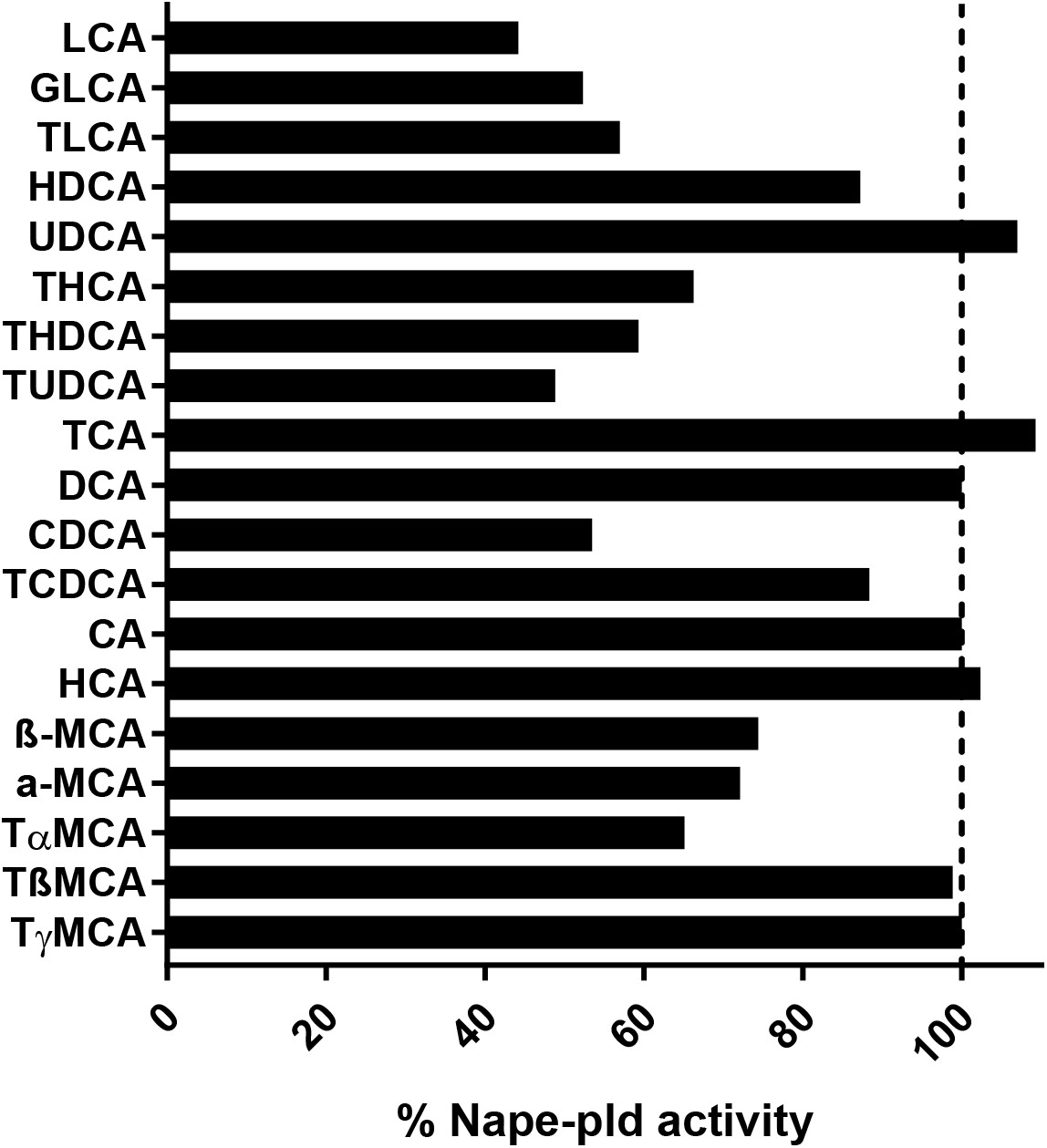
Effect of individual bile acid species (50 μM) on recombinant mouse Nape-pld activity measured by hydrolysis of PED-A1. Each bar represents the average of two replicates normalized to the activity of the DMSO negative control. LCA: lithocholic acid, GLCA: glycolithocholic acid, TLCA: taurolithocholic acid, HDCA: hyodeoxycholic acid, UDCA, ursodeoxycholic acid, THCA: taurohyocholic acid, THDCA: taurohyodeoxycholic acid, TUDCA: tauroursodeoxycholid acid, TCA: taurocholic acid, DCA: deoxycholic acid, CDCA: chenodeoxycholic acid, TCDCA: taurochenodeoxycholic acid, CA: cholic acid, HCA: hyocholic acid, MCA: muricholic acid, TαMCA: tauro-α-muricholic acid.

We next screened 2,388 biologically active and structurally diverse compounds (the Spectrum Collection compound library) at 10 μM final concentration of each compound. For all compounds, a B score (41) for both initial rate of fluorescence change and steady-state fluorescence value was calculated. Compounds with a B score 3 standards deviation above (activators) or below (inhibitors) the mean (or 3 median absolute deviations from the median) were classified as potential hits after confirmation by manual inspection (See Figure S3 in Supporting Information). Structures known to potentially interfere with the assay due to fluorescence or quenching were eliminated from further study. Hits were then re-assayed to confirm the original findings. While all 14 of the potential inhibitors demonstrated reproducible inhibition in the confirmation assay, none of the 12 potential activator compounds demonstrated reproducible activation.

To determine if the reduction in fluorescence by any of the 14 potential inhibitor compounds was due to direct quenching of BODIPY fluorescence, rather than inhibition of Nape-pld, these compounds were incubated with BODIPY FL C5, the PLA1 cleavage product of PED-A1. For each hit compound, the fluorescence of BODIPY FL C5 after treatment with 10 μM compound for 30 min was compared to vehicle treatment. None of the 14 hit compounds altered fluorescence by more than 10% (See Figure S4 in Supporting Information) indicating they were bona fide inhibitors of Nape-pld. All 14 inhibitor compounds (Inh 1-14) were then purchased from commercial sources to validate the putative library compound, analyzed by mass spectrometry to confirm identity and purity, and concentration response curve (CRC) studies performed to establish potency. The IC_50_ of each of these compounds ranged from 1.6 μM to 19.1 μM, so that each was potentially more potent in vitro than ARN19874 (Table 1) (37).

The effects of Inh 1 to 14 on cell viability were tested in HEK293 cells. The HEK293 cell line was chosen because it has been extensively used as a model for transfection and inhibition studies. Cytotoxicity of each compound was tested at a concentration 5 times that of their calculated IC_50_ in Table 1 (5xIC_50_). At these concentrations, six of the fourteen compounds gave cell viability >70%: Inh 1 (hexachlorophene), Inh 2 (bithionol), Inh 4 (ebselen), Inh 7 (estradiol valerate), Inh 12 (theaflavin mongallates), and Inh 14 (dioxybenzone) (Figure 3).

**Table 1:**
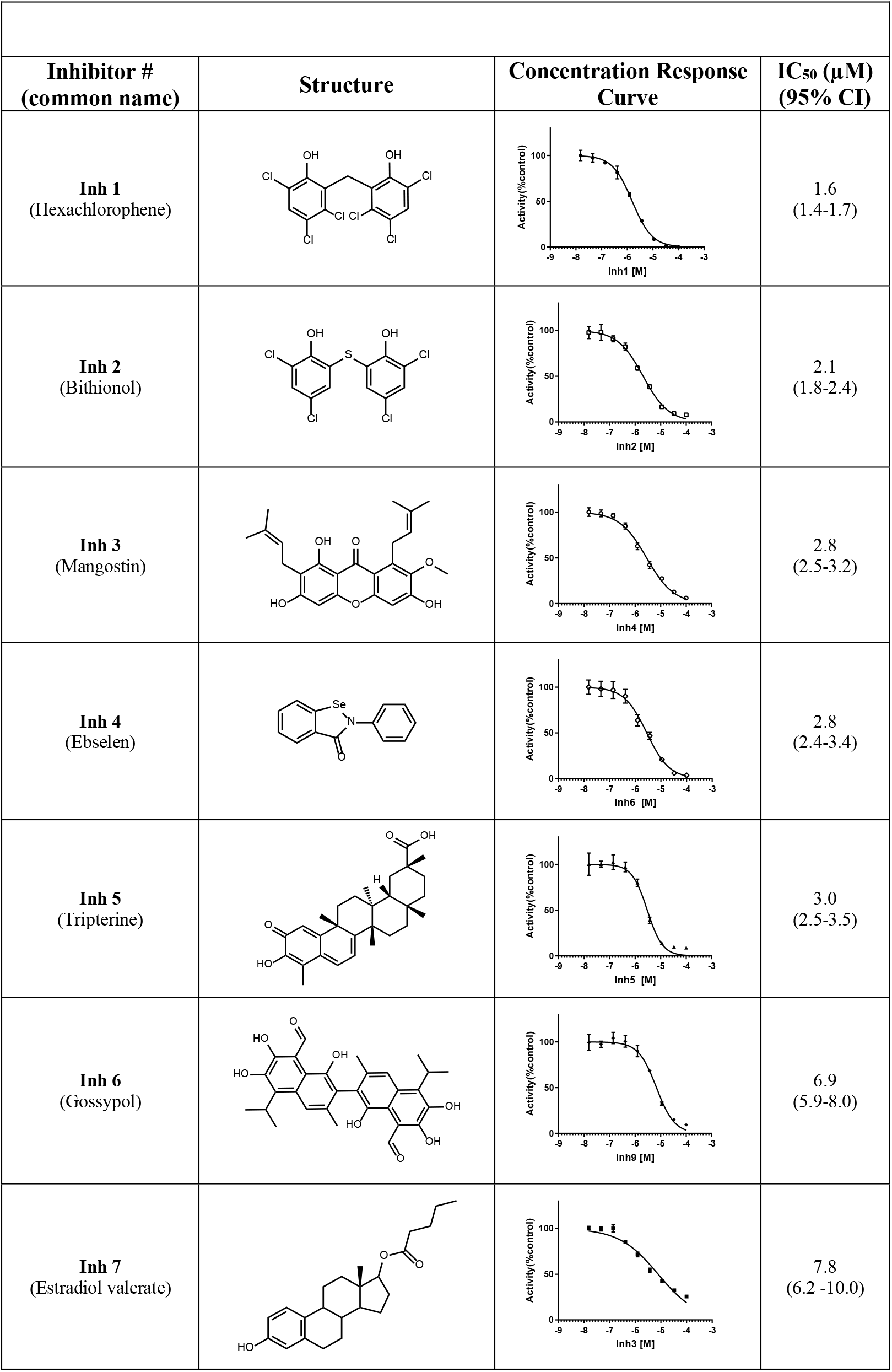
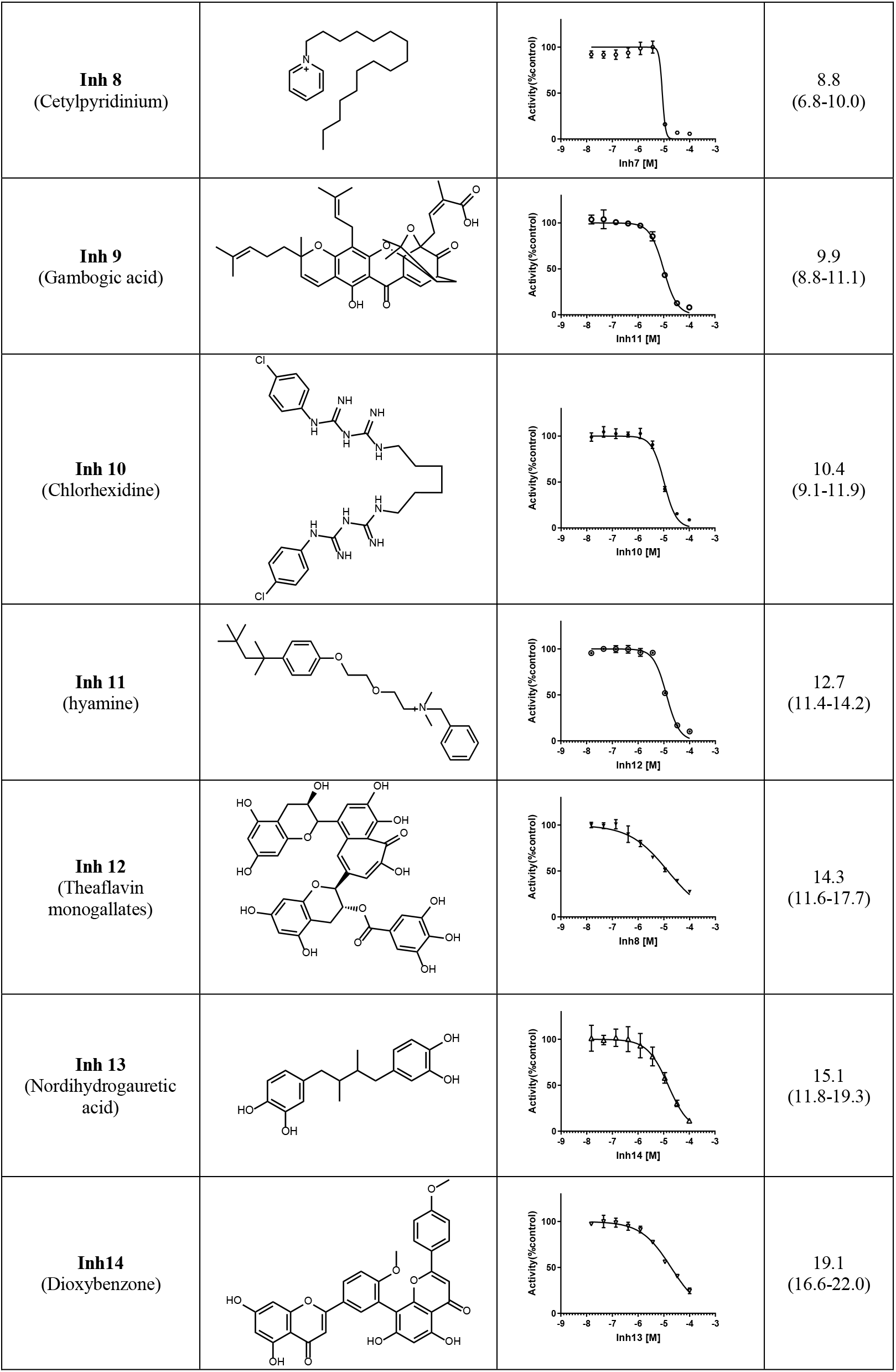
compounds identified from HTS screen of Spectrum Collection

**Figure 3.**
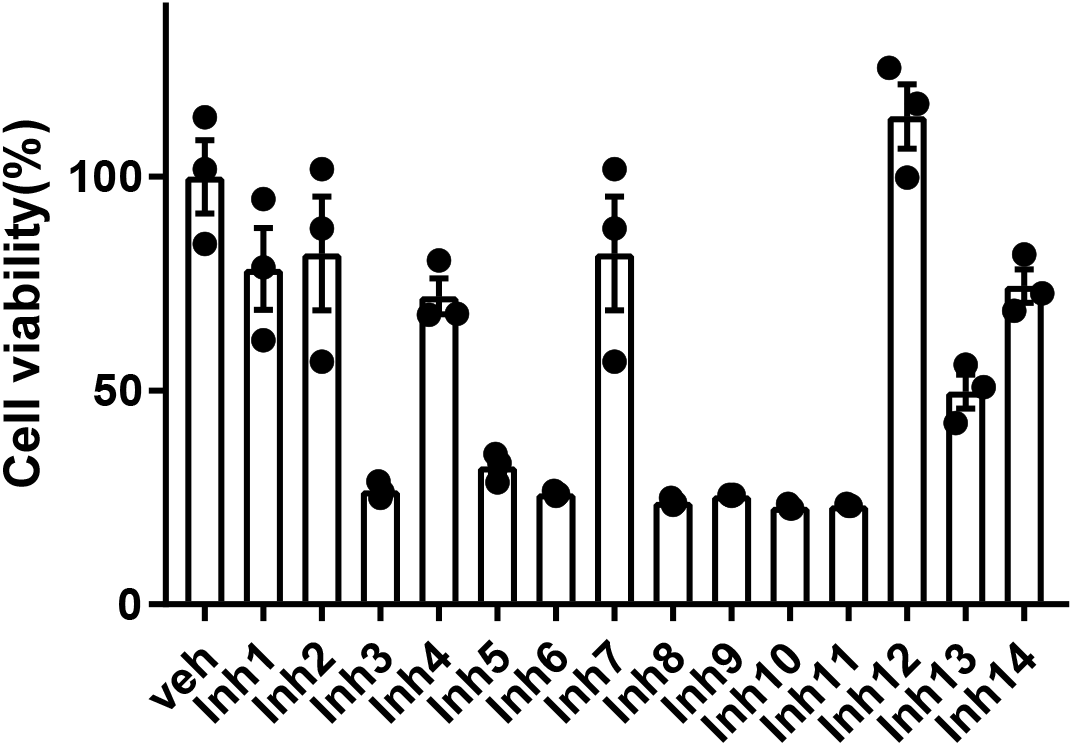
Effect of inhibitors on cell viability. HEK293 cells were incubated for 24 h with a concentration of inhibitor representing 5x the IC_50_ for recombinant Nape-pld for that compound. Resulting cell viability was measured by MTT assay and normalized to that of vehicle treated cells. Three replicate wells of 24-well plate were used for each compound. Mean±SEM shown for each compound.

Our finding that estradiol valerate (Inh 7, IC_50_ 7.8 μM) inhibited Nape-pld activity is of interest both because it shares significant structural homology to bile acids such as LCA and because estradiol valerate is a widely used birth control agent due to its potent estrogen receptor agonist activity. While this finding suggests the possibility that medicinal chemistry could be used to modify estradiol valerate to improve potency and reduce estrogen receptor activity, it is unclear what further modifications of this compound could be undertaken to this end. Several closely related estradiol analogs, including estradiol, estradiol diacetate, estradiol cypionate, and estratriol, were tested in the original screen of 2,388 compounds, but did not show significant inhibitory activity at 10 μM. Therefore, development of estradiol valerate was not pursued further.

Our two most potent inhibitors, Inh 1 (hexachlorophene) and Inh 2 (bithionol), share a symmetrically substituted dichlorophene structure, with Inh 2 having a thioether linker rather than an alkane linker, as well as lacking the 5/5’-chloro groups of Inh 1. Since Nape-pld acts as a homodimer, the potency of these symmetrical compounds may arise from acting at the dimer interface (e.g. by binding similar residues on both Nape-pld monomers). We used commercially available dichlorophene analogs to carry out limited structure activity relationship studies. We first tested whether the two hydroxyl groups contributed to inhibition. Inh 15, where chloro-groups replace the hydroxyl groups, had no inhibitory activity (Table 2). Inh 16 (chlorophene), with a hydroxyl and a chloro-group on only one of the two aromatic rings had only very weak inhibitory activity. Inh 17 (dichlorophen) and Inh 18 (fenticlor) have a hydroxyl and chloro-group on both aromatic rings and both of these compounds were about 10-fold less potent than Inh 1 and Inh 2. Inh 19, where both hydroxyl groups of Inh 17 are replaced with methoxy groups, gave no measurable inhibitory activity, supporting the need for symmetrical hydroxy groups. Symmetrical addition of methyl (Inh 20), chloro- (Inh 21), or bromo-groups (Inh 22) at the 2- and 2’-position to Inh 17 resulted in less, equal, or more potent inhibition of Nape-pld than for Inh 17, suggesting that addition of strong electronegative groups at this position enhances inhibitory activity. Addition of a trichloromethane group to the carbon bridging the two aromatic rings of Inh 17 (Inh 23) increased potency about 10-fold. Inh 24, where the 1-/1’-hydroxy groups of Inh 17 are at the 3-/3’-position instead, and there is dimethyl substitution of the carbon bridging the two aromatic rings, is about 3-fold less potent than Inh 17. Inh 25 (triclosan) also showed about 3-fold less potency than Inh 17, while Inh 26 (triclocarban) showed no inhibitory activity. Together, these results suggest that symmetrical halide substitution of the two phenolic groups is critical for potent inhibition. Although Inh 22 and Inh 23 showed similar potency of inhibition of recombinant Nape-pld as Inh 1 and Inh 2, both of them showed significantly greater cytotoxicity in HEK293 cells (data not shown), so we pursued further mechanistic testing only with Inh 1 and 2.

**Table 2:**
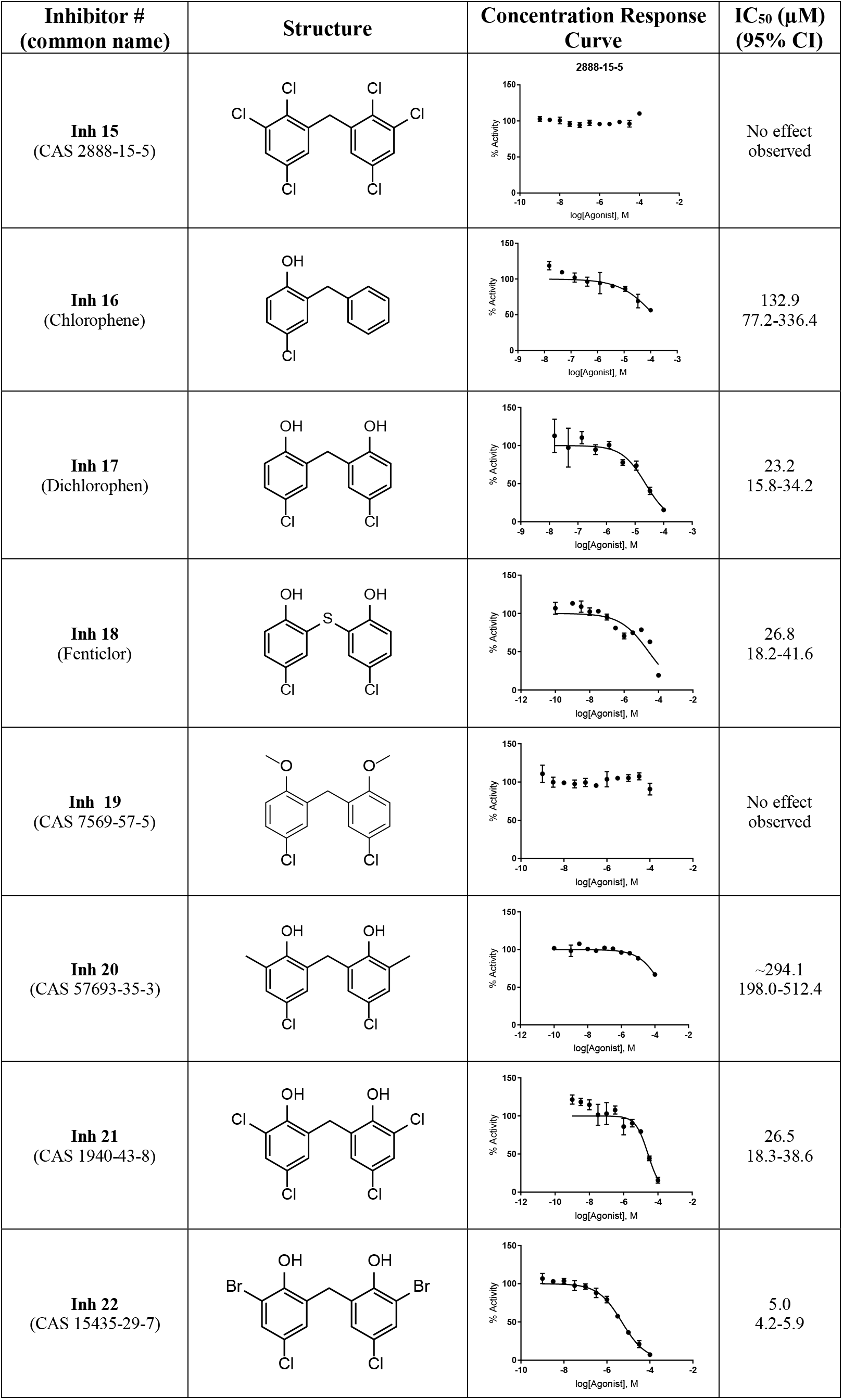
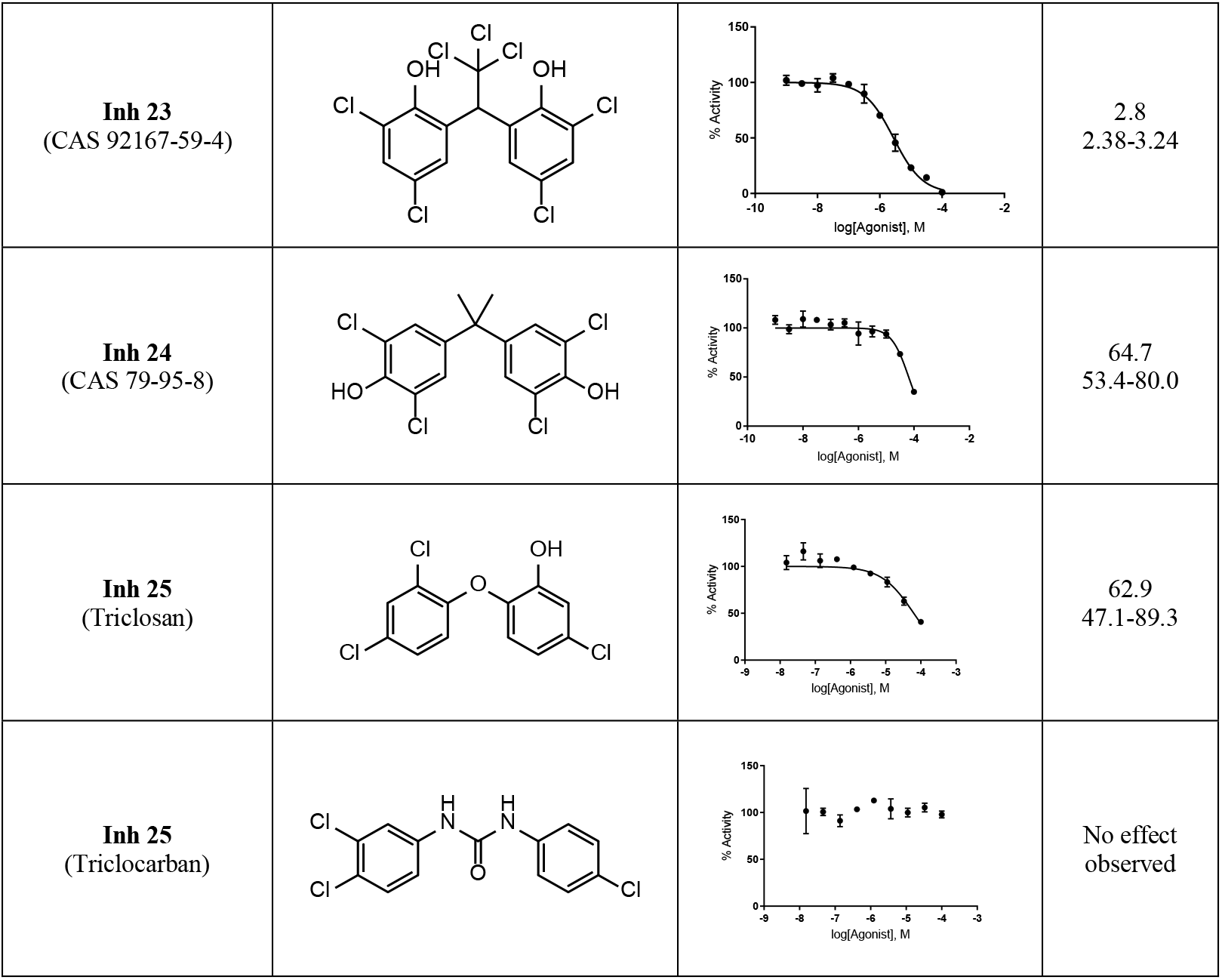
Structure activity relationships for substituted chlorophene analogs

Inhibition of recombinant Nape-pld with Inh 1 and Inh 2 altered V_max_ but not K_M_, consistent with a non-competitive mechanism of inhibition. Apparent KI for both Inh 1 and Inh 2 was ~2 μM (Figure 4A-B). Inhibition by Inh 1 was completely lost after 100-fold dilution of the enzyme inhibitor complex, indicating a non-covalent, highly reversible interaction (Figure 4C). In contrast, inhibition by Inh 2 persisted even after 100-fold dilution of the complex, indicating tight binding or potentially a covalent interaction. Thus, despite their chemical similarities, these two inhibitors appear to interact with Nape-pld by somewhat different mechanisms.

**Figure 4.**
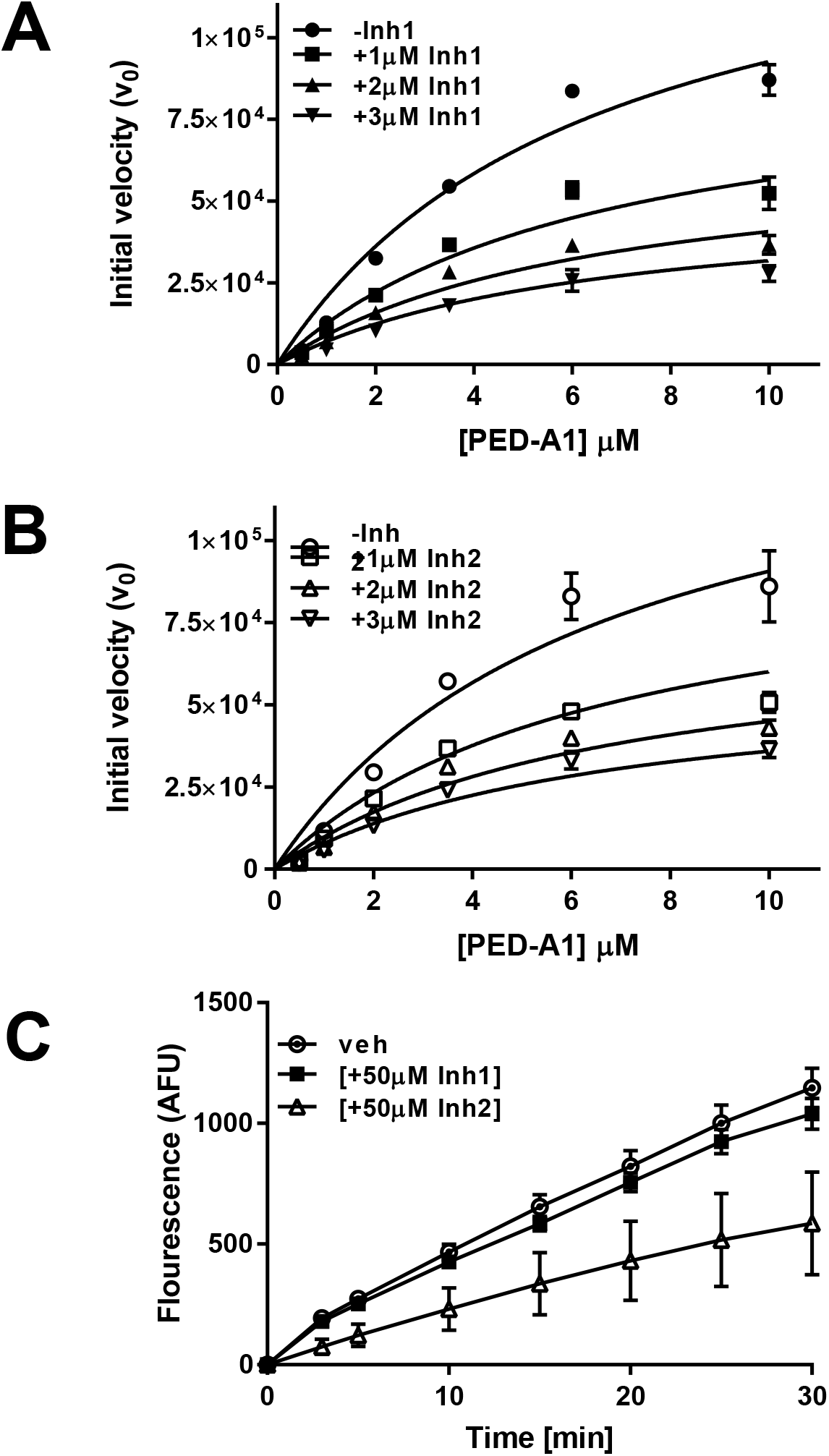
Mechanism of inhibition by Inh 1 and Inh 2. A) Recombinant Nape-pld was incubated with 0-3 mM Inh1 and then 0-10 mM PED-A1 added and the initial velocity of PED-A1 hydrolysis determined for each. (Mean ± SEM, n=3.) B) Recombinant Nape-pld was incubated with 0-3 mM Inh 2 and then 0-10 mM PED-A1 added and the initial velocity of PED-A1 hydrolysis determined for each. (Mean ± SEM, n=3.) C) A rapid dilution assay for Inh 1 and Inh 2. Recombinant Nape-pld was incubated with DMSO (veh) or 50 μM Inh 1 or Inh 2 for 1 h and then samples were diluted 100-fold with buffer immediately prior to addition of 4 μM PED-A1 and resulting fluorescence measured as arbitrary fluorescence units (AFU). (Mean ± SEM, n=9.)

Selectivity of the two inhibitors for Nape-pld versus other lipases was first examined using *Streptomyces chromofuscus* phospholipase D (ScPld), which is a broad-spectrum phospholipase D that catalyzes the hydrolysis of the headgroup of a variety of lipid substrates including phosphatidylcholine (PC), phosphatidylethanolamine (PE), NAPE, and other *N*-modified phospholipids (21–22,42). As with recombinant Nape-pld, incubation of ScPld with PED-A1 resulted in a rapid increase in fluorescence intensity. Both Inh 1 and Inh 2 inhibited ScPld activity, but only at concentrations ~100-fold higher (IC_50_ of 149 μM and >200 μM, respectively) than required to inhibit Nape-pld (Figure 5A). Potential inhibition of serum lipases was examined using 1-arachidonyl-thiolglycerol (1-AT) as a general lipase substrate, with released thiolglycerol quantified by fluorescence resulting from reaction with 7-diethylamino-3-(4’-maleimidylphenyl)-4-methylcoumarin. The IC_50_ for serum lipase activity was 110 μM and 46 μM for Inh 1 and Inh 2, respectively (Figure 5B). Because Nape-pld is a member of the zinc-dependent metalloenzyme family, we determined if Inh 1 and Inh 2 also inhibited other zinc metalloenzymes using carbonic anhydrase as a model enzyme. P-nitrophenylacetate (pNPA) was used as substrate to measure carbonic anhydrase activity. The IC_50_ for carbonic anhydrase activity was 107 μM and 178 μM for Inh 1 and Inh 2, respectively (Figure 5C). Thus, both Inh 1 and Inh 2 showed relatively selective inhibition of Nape-pld when compared to their effects on other lipases or zinc-dependent metalloenzymes.

**Figure 5.**
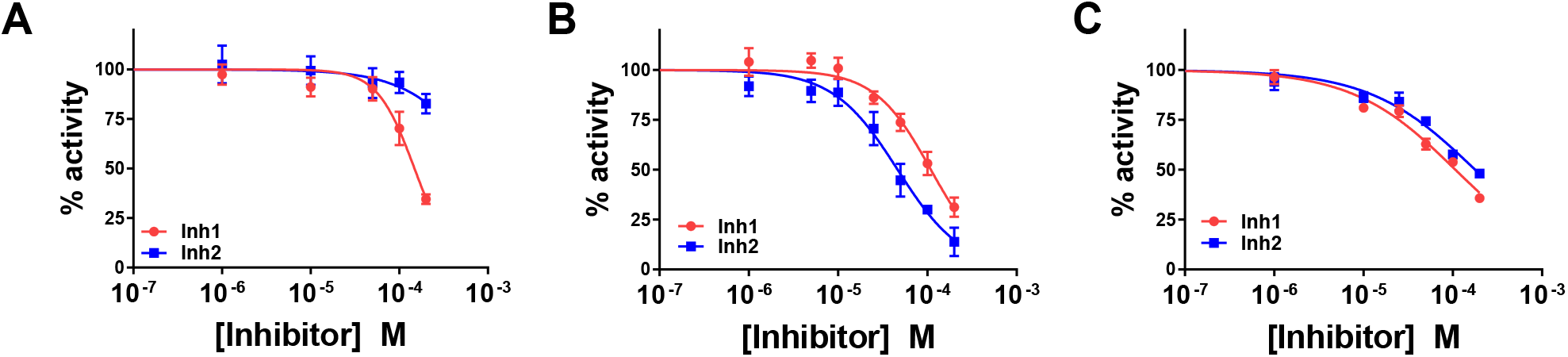
Selectivity of Inh 1 and Inh 2. A. Effect on *S. chromofuscus* phospholipase D activity. (Mean ± SEM, n=6.) B. Effect on serum lipase activity. (Mean ± SEM, n=9.) C. Effect on serum carbonic anhydrase activity. (Mean ± SEM, n=6.)

HEK293 endogenously express NAPE-PLD (37,43) and were therefore used to assess the ability of these inhibitors to inhibit the NAPE-PLD activity of intact cells. To determine the maximum concentration of Inh 1 and Inh 2 that could be used for cellular inhibition studies without cytotoxicity, HEK293 were treated with 0-100 μM inhibitor and cytotoxicity measured as before. Both inhibitors showed cytotoxicity at 100 μM (See Figure S5 in Supporting Information), so 20 μM was chosen as the maximum concentration for inhibition studies.HEK293 cells were treated with 0-20 μM of each inhibitor for 30 min and then NAPE-PLD activity measured by adding PED-A1. Inh 1 and Inh 2 inhibited NAPE-PLD activity with IC_50_ of 9.8 μM and 10.7 μM, respectively (Figure 6A-B).

**Figure 6.**
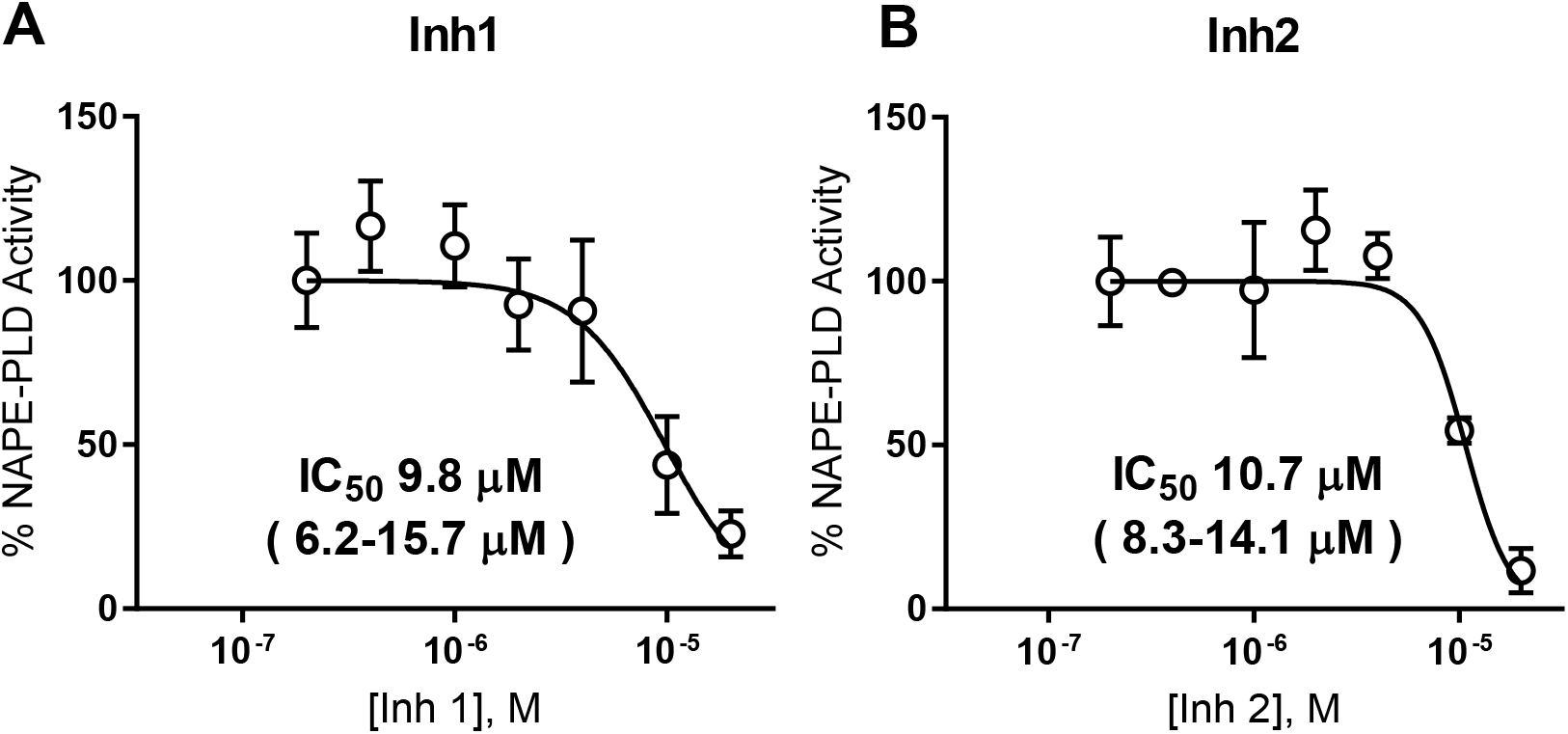
Inh 1 and Inh 2 inhibit the NAPE-PLD activity of HEK293 cells. 0-20 mM of Inh1 (A) or Inh 2 (B) were added to confluent wells of HEK293 cells plated in 96-well plates and incubated for 30 min. PED-A1 was then added and NAPE-PLD activity determined by change in fluorescence (ex/em 488/530nm) over 25 min and then normalized to control. Points represent mean ± SEM, n = 6 (3 wells on two replicate days). IC_50_ and 95% confidence interval calculated using log(inhibitor) vs. normalized response-variable slope analysis.

From these studies, we conclude that symmetrically substituted dichlorophene are potent inhibitors of NAPE-PLD in cultured cells and show significant selectivity for NAPE-PLD versus other tissue lipases. Both Inh 1 and Inh 2 can therefore be used for cultured cell experiments to test the effects of NAPE-PLD inhibition. Although both inhibitors are relatively selective for NAPE-PLD compared to other lipases, both are known to have other off-target effects not related to lipases, so care must be taken to ensure non-toxic concentrations are used as both inhibitors are known anti-helminth, antibacterial, and antifungal agents.

## Experimental procedures

### Reagents

Recombinant hexahistidine-tagged mouse Nape-pld was expressed and purified as described previously (43). N-((6-(2,4-DNP)Amino)Hexanoyl)-1-(BODIPY™ FL C5)-2-Hexyl-Sn-Glycero-3-Phosphoethanolamine (PED-A1) and BODIPY™ FL C5 were purchased from Thermofisher scientific, USA. Rabbit anti-NAPE-PLD primary antibody was obtained from Abcam. Human embryonic kidney 293 (HEK293) cells were purchased from American Type Culture Collection (Manassas, VA). The fourteen hit inhibitor compounds used for confirmation were purchased from MicroSource Discovery Systems, Inc. Dulbecco’s modified Eagle’s medium (DMEM medium), optiMEM (1X) reduced serum medium and heat-inactivated fetal bovine serum (HI-FBS) were from Gibco chemicals. Streptomyces chromofuscus PLD (ScPld), p-nitrophenylacetate, lithocholic acid, 2-acetazolamide and 3-(4,5-dimethylthiazol-2-yl)-2,5-diphenyltetrazolium bromide (MTT) were purchased from Millipore-Sigma. 1-Arachidonyl thioglycerol, N-octyl-β-D-glucoside (NOG) and 7-diethylamino-3-(4’-maleimidylphenyl)-4-methylcoumarin were obtained from Cayman Chemicals. Standard protein molecular weight marker was purchased from Invitrogen. Nickelnitrilotriacetic acid-agarose was from GE Healthcare. Deidentified, surplus human serum was provided by the Vanderbilt Clinical Pharmacology Phlebotomy Core. The Spectrum Collection compound library was provided by the Vanderbilt University High Throughput Screening Core facility and was originally obtained from MicroSource (Gaylordville, CT). The collection includes a wide range of biologically active and structurally diverse compounds consisting of active drugs (50%), natural products (30%) and other bioactive components (20%).

### Initial Nape-pld activity assay

Recombinant mouse Nape-pld or boiled Nape-pld was diluted in assay buffer containing 50mM Tris-HCl, pH 8.0 and 1% NOG to a final concentration of 1.0 μM and incubated with 10 μM PED-A1. Fluorescence (ex/em 488/530nm, fixed bandwith 16 nm) was measured at 1-minute time points for 2 h in 96-well plate reader (Synergy H1 Hybrid Multi-Mode Reader, BioTeK).

### NAPE-PLD activity assay optimization

KM for PED-A1 hydrolysis by recombinant mouse Nape-pld was determined by enzyme kinetics. Briefly, Nape-pld was diluted in assay buffer to a final concentration of 4.5 μg mL-1 and enzyme kinetics was performed in a 384-well plate using 0– 20 μM PED-A1. Fluorescence (excitation 482±18 nm; emission 536±20 nm) was measured for 15 min using a Wavefront Panoptic kinetic imaging plate reader. Effect of varying concentration of NOG (0–1%) concentration were measured using final concentration of 4 μM PED-A1 and 4.5 μg mL-1 Nape-pld. For the DMSO tolerance test, DMSO (35 nL–900 nL) was transferred onto an empty 384-well plate using an Echo liquid handler. Then, 30μL of 4.5 μg mL-1 enzyme diluted in assay buffer containing 0.4% NOG (w/v) was dispensed into the DMSO-containing plate using a Bravo liquid handler robot, followed by 5 μL PED-A1 (final concentration 4 μM) and fluorescence kinetics (ex/em 482/536 nm) measured for 15 min. The consistency of replicates across the 384-plate was determined using the final optimized concentrations of DMSO (1.6%, v/v), NOG (0.4%, w/v), and 4.5 μg mL-1 Nape-pld, with replicates of the NAPE-PLD inhibitor, lithocholic acid (LCA), dispersed in checkerboard pattern across the plate (“Checkerboard Assay”), and then 5 μL PED-A1 (4 μM final concentration) added. The Z-prime factor for LCA inhibition across the plate was calculated as Z=1-[3(X + X’)/(Y – Y’)], where Y and Y’ are the mean values of PED-A1 produced fluorescence in the absence and presence, respectively, of LCA as a control inhibitor at initial rate of reaction. X and X’ are standard deviation of PED-A1 produced fluorescence.

### High-throughput screen

Screening was performed on 2,388 compounds. 35 nL of 10 mM test compounds and 140 nL of 100% DMSO were combined in black-walled, clear bottom 384-well plates using an Echo Acoustic liquid handler, resulting in a final compound concentration of 10 μM for the primary screen. 175 nL of 20 mM LCA or 175 nL of DMSO were added to control wells as inhibited and uninhibited signal controls, respectively. 30 μL of 4.5 μg mL-1 Nape-pld enzyme in buffer solution with 0.4% NOG (w/v) was dispensed in each well using an Agilent Bravo robotic liquid handler. Plates were incubated at 37°C for 1hr. The assay was initiated by the Panoptic instrument’s internal Bravo robotic liquid handler when it dispensed 5 μL of PED-A1 (4 μM final) in buffer solution (1.6% DMSO final, v/v) to each well. Changes in fluorescence (ex/em 482/536nm) were measured for 15 min. Initial slope measurements used slope from t = 30-100 sec.

### Concentration-response curve

The IC_50_ for each hit compound was determined using similar conditions as for screening except that concentration of inhibitor was varied from 0 to 200 μM. The averaged initial slope measurements for NAPE-PLD activity in the absence of inhibitor were set as 100% activity and the normalized % activity calculated by dividing the initial slope observed with each concentration of inhibitor by this value. Fitting of the response curve and calculation of the IC_50_ and 95% confidence interval was performed using log(inhibitor) vs. normalized response-variable slope analysis in GraphPad Prism version 7.04.

### Rapid dilution assay

This assay was performed using the methods of Castellani et al (37). Briefly, 81 μg mL-1 recombinant Nape-pld was pre-incubated with Inh 1 or Inh 2 at 50 μM for 1 h and then samples were diluted 100-fold with 50 mM Tris-HCl (pH=8.0) immediately prior to addition of 4 μM PED-A1 and fluorescence measured at 1-min intervals for 30 min.

### Cytotoxicity

Effect of compounds on viability of HEK293 cells were determined using MTT assay to measure changes in cellular redox state. Briefly, 10×10^4^ HEK293 cells/well were seeded in 24-well plates for 24 h. Cells were treated with a concentration of each compound that represented 500% of the IC_50_ concentration determined in the CRC assay (5×IC_50_) prepared in opti-MEM reduced serum media containing 1% HI-FBS for 24 h. After treatment, media was removed, and 300 μL of 0.5 mg mL-1 MTT solution was added. After 3 h, the media was removed, the purple crystals dissolved in 100 μL of DMSO, and 80 μL transferred to 96-well plate, and absorbance measured at 590 nm. Percent viability was normalized to cells treated with vehicle (DMSO) only.

### Cellular NAPE-PLD activity

HEK293 cells in DMEM buffer containing 4.5 g L-1 glucose, 4 mM L-glutamine, 1 mM sodium pyruvate, phenol red and supplemented with 10% HI-FBS were seeded at ~20,000 cells/well in tissue culture-treated, clear-bottom, black-walled, 96-well plates. After reaching ~95% confluency (48hr), the medium was removed and replaced with similar DMEM buffer (70 μL per well) except without phenol red or HI-FBS. 5 μL of inhibitor or vehicle solutions were then added to each well to generate final concentrations of inhibitors of 0.2–20 μM. 30 min after addition of inhibitors, 5 μL of 56 μM PED-A1 was added to each well (3.5 μM final concentration), and fluorescence signal (ex/em 488/530nm) was recorded every minute for 25 min in Synergy H1 plate reader. NAPE-PLD activity was measured as the increase in fluorescence from time = 1 to 25 min and normalized to average activity found in absence of inhibitor. For each concentration of inhibitor, 3 replicate wells on two separate days were measured (n=6 wells total) and mean ± SEM determined.

### Counter-screens and specificity screens

All assays were carried out in 96-well plates. To assess effect of compounds on BODIPY fluorescence, 4 μM of BODIPY-FL C5 was incubated with or without 10 μM of each compound in total 80μL of 50 mM Tris-HCl (pH=8.0) and fluorescence (ex/em 482/536nm) measured for 30min. To assess effects of compound on *S. chromofuscus* phospholipase D (ScPld) activity, 0.1 μM ScPLD was incubated with 0–200 μM of Inh 1 or Inh 2 in 70 μL of 50 mM Tris-HCl (pH=8.0) for 1 h, PED-A1 added to 4 μM, and fluorescence (ex/em 488/530nm) measured by Synergy H1 plate reader for 15 min. For assays assessing the effect of compounds on serum enzymes, deidentified surplus human serum was obtained from the Vanderbilt Clinical Pharmacology Phlebotomy Core and diluted 1:9 (i.e. 10-fold dilution factor) using 50 mM Tris-HCl (pH=8.0) to generate a diluted serum stock solution. Pan-lipase activity was measured using a lipase assay kit (Cayman Chemical, 700640) which uses 1-arachidonyl-thiolglycerol (1-AT) as substrate (final concentration 23 μM) and the released thioglycerol detected using thiol detection agent (7-dethylamino-3-(4’-Maleimidylphenyl)-4-Methylcoumarin, final concentration 50 μM). Recombinant mouse Nape-pld (4.5 mg L-1) did not hydrolyze 1-AT to any appreciable extent, while the bovine milk lipoprotein lipase supplied in the assay kit (diluted according to manufacturers recommendation of 10 μl supplied stock diluted with 140 μl assay buffer) gave robust 1-AT hydrolysis (See Figure S6A in Supporting Information). To determine the K_m_ for 1-AT hydrolysis activity in diluted serum, 1-AT concentration was varied from 0-90 μM. This gave an estimated K_m_ of 21.5± 3.4 μM (See Figure S6B in Supporting Information). To measure effect of inhibitors on serum lipase activity, 5 μL of Inh 1, Inh 2, or vehicle (DMSO) (final inhibitor concentration 0–200 μM) was added to each well containing 60 μL of 50 mM Tris-HCl (pH=8.0) and 10 μL of diluted serum stock solution and incubated together for 30 min. 5 μL of a solution containing 1-arachidonyl-thiolglycerol and thiol detection agent (7-dethylamino-3-(4’-Maleimidylphenyl)-4-Methylcoumarin), (final concentration 23 μM and 50 μM, respectively) was then added and change in fluorescence (ex/em 380/515 nm) monitored using Synergy H1 plate reader for 30 min. 100% activity was set as the fluorescence detected for vehicle only. Serum carbonic anhydrase activity was measured using a carbonic anhydrase assay kit (Biovision Inc., K472) which uses p-nitrophenylacetate (1 mM) as substrate and the released p-nitrophenol measured by absorbance at 405 nm. The positive control carbonic anhydrase provided in the assay kit robustly hydrolyzed p-nitrophenylacetate while recombinant Nape-pld (4.5 mg L-1) did not to any appreciable extent (See Figure S7A in Supporting Information). Varying p-nitrophenylacetate concentration from 0-5 mM gave an estimated KM of 2 mM using diluted serum (See Figure S7B in Supporting Information). A known carbonic anhydrase inhibitor, acetazolamide (1 mM), robustly inhibited p-nitrophenylacetate hydrolysis activity of diluted serum (See Figure S7C in Supporting Information). To assess the effect of inhibitors on serum carbonic anhydrase activity, 5 μL of Inh 1, Inh 2, or vehicle (DMSO) (final inhibitor concentration 0–200μM) was added to 60 μL of 50 mM Tris-HCl (pH=8.0) and 10 μL of diluted serum stock solution per well in 96-well plate and incubated together for 30 min. 5 μL of p-nitrophenylacetate (1 mM final concentration) was then added and hydrolysis by carbonic anhydrase measured as change in absorbance at 405 nm in Synergy H1 plate reader.

### Statistical analysis

In all reactions, the IC_50_ was calculated using log(inhibitor) vs. normalized response-variable slope analysis in GraphPad Prism version 7.04.

## Supporting Information

Supplemental figures (Figures S1-S7).

## Acknowledgments

This work was supported by a Discovery Grant from the Vanderbilt Diabetes Center (SSD) and by the NIH (DK105847, CRF). The WaveFront Biosciences Panoptic kinetic imaging plate reader is housed and managed within the Vanderbilt High-Throughput Screening Core Facility, an institutionally supported core, and was funded by NIH Shared Instrumentation Grant 1S10OD021734. The Microsource Spectrum Collection was distributed by the Vanderbilt High-Throughput Screening Core Facility with assistance from Ms. Corbin Whitwell. The HTS Core receives support from the Vanderbilt Institute of Chemical Biology and the Vanderbilt Ingram Cancer Center, which in turn receives support from NIH (P30 CA68485).

